# The String Decomposition Problem and its Applications to Centromere Assembly

**DOI:** 10.1101/2019.12.26.888685

**Authors:** Tatiana Dvorkina, Andrey V. Bzikadze, Pavel A. Pevzner

## Abstract

Recent attempts to assemble long tandem repeats (such as multi-megabase long centromeres) faced the challenge of accurate translation of long error-prone reads from the nucleotide alphabet into the alphabet of repeat *units*. Centromeres represent a particularly complex type of *nested tandem repeats*, where each unit is itself a repeat formed by chromosome-specific *monomers* (a repeat within repeat). Given a set of monomers forming a specific centromere, translation of a read into monomers is modeled as the String Decomposition Problem, finding a concatenate of monomers with the highest-scoring sequence alignment to a given read. We developed a StringDecomposer algorithm for solving this problem, benchmarked it on the set of reads generated by the Telomere-to-Telomere consortium, and identified a novel (rare) monomer that extends the set of twelve X-chromosome specific monomers identified more than three decades ago. The accurate translation of each read into a monomer alphabet turns centromere assembly into a more tractable problem than the notoriously difficult problem of assembling centromeres in the nucleotide alphabet. Our identification of a novel monomer emphasizes the importance of careful identification of all (even rare) monomers for follow-up centromere assembly efforts.

## Introduction

Recent advancements in long read sequencing technologies, such as Pacific Biosciences (PB) and Oxford Nanopore Technologies (ONT), led to a substantial increase in the contiguity of genome assemblies (Koren et al., 2017; Kolmogorov et al., 2019; Ruan and Li, 2019; Shafin et al., 2019) and opened a possibility to resolve *extra-long tandem repeats* (*ETRs*), the problem that was viewed as intractable until recently. Assembling ETRs is important since variations in ETR have been linked to cancer and infertility (Barra and Fachinetti, 2018; Black and Giunta, 2018; Ferreira et al., 2015; Giunta and Funabiki, 2017; Miga et al., 2019; Smurova and De Wulf, 2018; Zhu et al., 2018). ETR sequencing is also important for addressing open problems about centromere evolution (Alkan et al., 2007; Lower et al., 2018; Shepelev et al., 2009, Henikoff et al., 2015).

The initial attempts to assemble ETRs (Jain et al., 2018; Bzikadze and Pevzner, 2019; Miga et al., 2019) revealed the importance of the *String Decomposition Problem*, partitioning an ETR (or an error-prone long read sampled from an ETR) into repetitive *units* forming these repeats. Although no existing tool explicitly addresses the String Decomposition Problem, Tandem Repeats Finder (Benson, 1999), PERCON (Kazakov et al., 2003), Alpha-CENTAURI (Sevim et al., 2016), and the Noise-Cancelling Repeat Finder (Harris et al., 2019) address related problems. Although these tools can be adapted for string decomposition, they often result in limited accuracy in the case of *nested tandem repeats*, such as centromeres and rDNA arrays.

Centromeres are the longest tandem repeats in the human genome that are formed by units repeating hundreds or even thousands of times with extensive variations in copy numbers in the human population and limited nucleotide-level variations. Each such unit (referred to as *high-order repeat* or *HOR*) usually represents a tandem repeat formed by smaller building blocks (referred to as *alpha-satellites* or *monomers*), thus forming a nested tandem repeat, i.e., a repeat within another repeat.

The alpha satellite repeat family occupies around 3% of the human genome (Hayden et al., 2013). Each monomer is of length ∼171bp and each HOR is formed by multiple monomers. For example, the vast majority of HORs on human X centromere (referred to as cenX) consist of twelve monomers. Although different HOR units on cenX are highly similar (95-100% sequence identity), the twelve monomers forming each HOR are rather diverged (50-90% sequence identity). In addition to standard 12-monomer HORs, some units on cenX have non-canonical monomer structure: 35 out of 1510 units are formed by smaller or larger number of monomers than the canonical 12-mer unit (Bzikadze and Pevzner, 2019). Moreover, two 12-mer units on cenX represent a non-canonical order of monomers. The tandem repeat structure of human centromeres may be interrupted by retrotransposon insertions (for example, cenX has a single insertion of a LINE element).

Partitioning of long error-prone reads into units and monomers is critically important for centromere assembly (Bzikadze and Pevzner, 2019; Miga et al., 2019). For example, centroFlye (Bzikadze and Pevzner, 2019) requires a translation of each read in the nucleotide alphabet into a *monoread* in the monomer alphabet. Unless these monoreads are extremely accurate (e.g., less than 0.5% error rate), the centroFlye assembly fails. However, the existing tools for analyzing tandem repeats (Benson, 1999; Harris et al., 2019; Sevim et al., 2016) generate monoreads with higher error rates and thus do not provide an adequate solution of the String Decomposition Problem.

Here we present a StringDecomposer (SD) algorithm that takes the set of monomers and a long error-prone read (or a genomic segment) and partitions this read into distinct monomers. The accurate translation of each read from a nucleotide alphabet into a monomer alphabet opens a possibility to assemble the reads in the monomer alphabet, a more tractable problem than the notoriously difficult problem of assembling ETRs in the nucleotide alphabet.

## Methods

### String Decomposition Problem

Given a string *R* (corresponding to a read or an assembly of a centromere) and a set of strings *Blocks* (each *block* from *Blocks* corresponds to an alpha-satellite), the goal of the String Decomposition Problem is to represent *R* in the alphabet of blocks. We define a *chain* as an arbitrary concatenate of blocks, and an *optimal chain* for *R* as a chain that has the highest-scoring global alignment against *R* among all possible chains. The String Decomposition Problem is to find an optimal chain for *R*.

### String Decomposition Graph

Given a block *b* and a string *R*, their standard *alignment graph* consists of all vertices (*i,j*) for 0 ≤ *i* ≤ |*b*|, and 0 ≤ *j* ≤ |*R*|, where |·| stands for the length of a string. A vertex (*i,j*) is connected to vertices (*i+1,j*), (*i,j+1*), and (*i+1,j+1*) representing insertions, deletions, and matches/mismatches, respectively.

Informally, given a set of *t* blocks and a string *R*, their *String Decomposition Graph* consists of *t* standard alignment graphs that are “glued together” by their 0-th row (Figure 1). We index vertices in this graph as (*b,i,j*) using three indices (*b* represents a block, *i* represents a position in the block *b*, and *j* represents a position in string *R*) except for vertices in the 0-th row that are indexed simply as (0,*j*) since all blocks share the same “glued” 0-th row. Additionally, we add edges that connect each vertex (*b,|b|,j*) in the last row (for each block *b*) with vertex (0,*j*) in the 0-th row.

**Figure 1.**
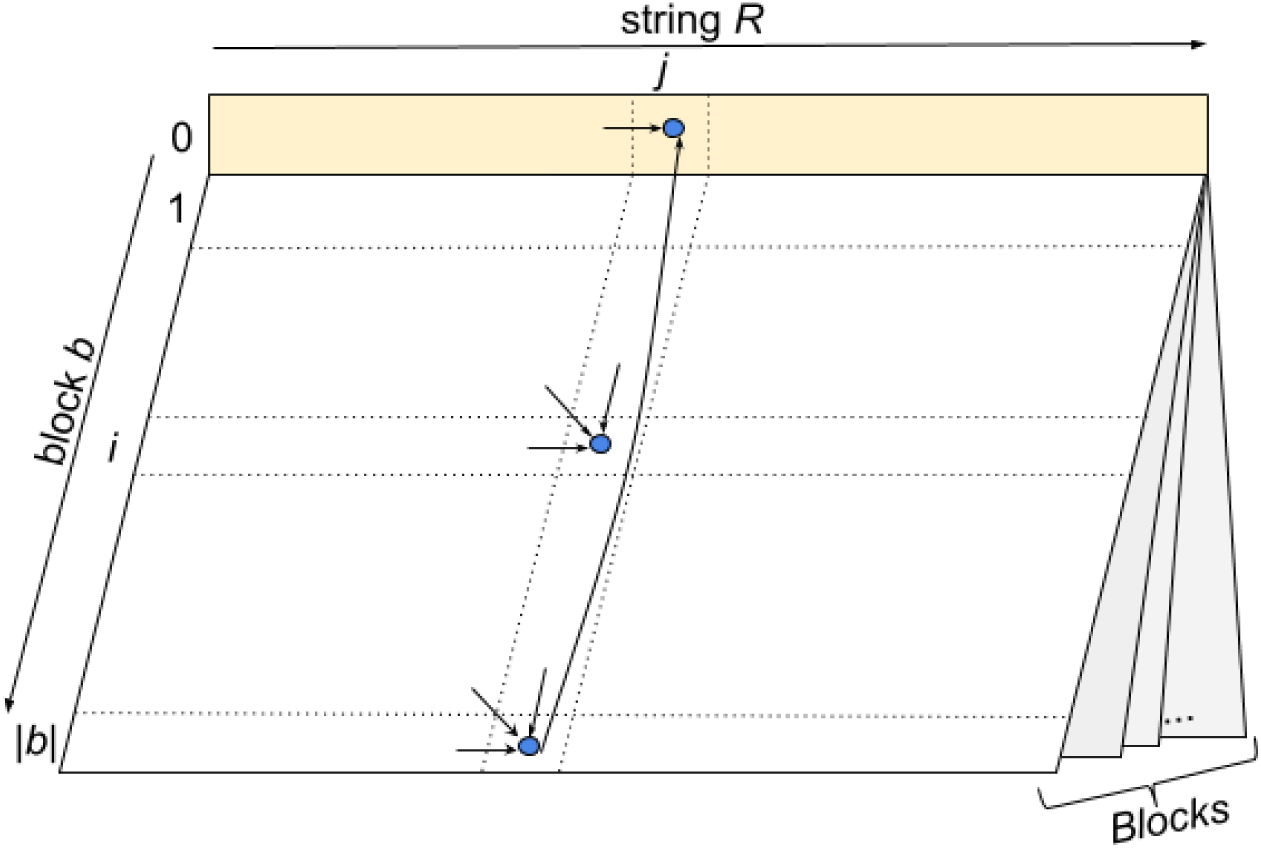
The String Decomposition Graph, represented as a “book” where each page corresponds to an alignment matrix for a single block, and pages are “glued” together by their 0-th row. Arrows represent edges of the String Decomposition Graph.

Formally, the String Decomposition Graph graph is constructed on all vertices (*b,i,j*), where *b* is a block, 0 ≤ *i* ≤ |*b*|, and 0 ≤ *j* ≤ |*R*| under the assumption that, for each *j*, all vertices (*b*,0,*j*) form a single vertex (0,*j*). A vertex (*b,i,j*) is connected to vertices (*b,i+1,j*), (*b,i,j+1*), and (*b,i+1,j+1*) representing insertions, deletions, and matches/mismatches, respectively (as in the standard alignment graph), and scored as –*δ* for insertions/deletions (indels), -*σ* for mismatches, and +1 for matches. Additionally, vertex (*b*, |*b*|, *j*) is connected by a zero-weight *block-switching edge* to a vertex (0, *j*) for each block *b* from the block-set. Although the String Decomposition Graph has directed cycles, the longest path in this graph is well-defined since all directed cycles have negative weights. We refer to the vertex (0,0) as the *source* and to each vertex (*b*,|*b*|,|*R*|) as a *sink* of the String Decomposition Graph.

### StringDecomposer algorithm for solving the String Decomposition Problem

The *i*-*prefix* of a block is defined as the string formed by its first *i* symbols. Given a block *b*, we define a (*b,i*)-*chain* as a chain complemented by the *i*-prefix of *b* in the end (note, that multiple (*b,i*)-chains exist). A dynamic programming algorithm for solving the String Decomposition Problem is based on computing a variable

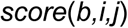

defined as the score of an optimal global alignment between all possible *(b,i)*-chains and the *j*-prefix of *R*. We assume that the alignment is scored using the following parameters: –*δ* for indel penalties, *-σ* for mismatch penalty, and +1 for matches. Similarly to the *fitting alignment* (Gusfield, 1997), the variable *score*(*b,i,j*) is initialized as *score*(*b,0,j*)*=*0 for all *b* and *j* and *score*(*b,i,0*)*=-i*\**δ* for all *i*.

To compute *score*(*b,i,j*), the StringDecomposer (SD) algorithm uses dynamic programming to compute the length of a longest path to the vertex *(b,i,j)* in the String Decomposition Graph. The solution of the String Decomposition Problem is given by the length of a longest path from the source to one of the sinks:

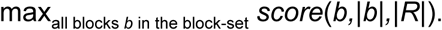

A block *b* that maximizes this score represents the last block in an optimal chain. Other blocks in this chain are inferred by backtracking from the sink (*b*,|*b*|,|*R*|) to the source (0,0) and are defined by the block-switching edges in the backtracking path.

The running time of the SD algorithm for solving the String Decomposition Problem for a string *R* and a block-set *Blocks* is O(|*R*|*\**(l*ength*(*Blocks*) *+ number*(*Blocks*)^*2*^)), where *length*(*Blocks*) and *number*(*Blocks*) is the total length of all blocks and the number of blocks, respectively (see Appendix “<u>SD implementation details</u>”). The memory footprint is O(|*R*|*\*length*(*Blocks*)).

### Transformation from the nucleotide alphabet to the block alphabet

Given a string *R* and a block-set *Blocks*, SD generates an optimal chain for *R.* For each block *b* in an optimal chain, SD outputs the starting and ending positions of the alignment of this block in *R.* We denote the substring of *R* spanning these positions as *R(b*), construct an alignment between the block *b* and *R*(*b*), and compute the percent identity of this alignment referred to as *IdentityR(b)*. A block *b* in an optimal decomposition of a string *R* is called *reliable* (*unreliable*) if *IdentityR(b)* exceeds (does not exceed) the threshold *MinIdentity* with default value *MinIdentity* = 69% (see Appendix “<u>SD implementation details</u>”). We substitute all unreliable blocks in the string decomposition by the *gap symbol* “?”, resulting in a translation of the string *R* into the *extended block alphabet* that consists of all blocks and the gap symbol (see Appendix “<u>Processing gaps in the monomer alignment</u>”). The translated sequence is referred to as *translation*(*R)*. If the string *R* is a centromeric read *Read* and blocks represent monomers, we refer to the *translation*(*Read)* as *mono(Read).*

### Existing approaches to solving the String Decomposition Problem

Although no existing tool explicitly addresses the String Decomposition Problem, Minimap2 (Li, 2018), PERCON (Kazakov et al., 2003), Tandem Repeats Finder (Benson, 1999), Alpha-CENTAURI (Sevim et al., 2016), and Noise-Cancelling Repeat Finder (Harris et al., 2019) tools address related problems.

Harris et al., 2019 demonstrated that the performance of general purpose sequence aligners such as Minimap2 (Li, 2018) deteriorates in highly repetitive regions, making them not suitable for solving the String Decomposition Problem.

PERCON (Kazakov et al., 2003) is a fast heuristic algorithm for solving the String Decomposition Problem that compares octanucleotide content of a potential monomer sequence with all known monomers. Unfortunately, PERCON is difficult to benchmark since it was implemented only for Windows OS.

Tandem Repeats Finder (TRF) is a popular *de novo* tandem repeat finder, that does not require specifying either the monomer or its size as an input (Benson, 1999). Given a string containing a tandem repeat with unknown monomers, it reports a consensus of these monomers and the location of each monomer in the repeat. TRF identifies monomers of length up to 2000 bp and is able to identify alpha satellites. However, the output of TRF can not be directly converted into a chain because TRF neither attempts to identify different monomer classes from the input string itself, not accepts monomer classes as an input. A drawback of this approach is that it often reports a cyclic shift of a consensus monomer and locations of this cyclic shift in the repeat, making it difficult to benchmark TRF against other string decomposition tools (see Appendix “<u>Benchmarking string decomposition tools</u>”).

The Noise-Cancelling Repeat Finder (NCRF; Harris et al., 2019) takes a consensus HOR as an input and partitions an error-prone read into HORs. It performs well if the entire read is formed by *canonical* HORs (formed by the same order of monomers as the consensus HOR). However, it generates a suboptimal and somewhat arbitrary partitioning into HORs when a read contains non-canonical HORs or retrotransposons. Since NCRF was not designed for decomposing reads into distinct monomers, it is difficult to benchmark it against monomer-finding tools (see Appendix “<u>Monomer-free benchmarking</u>”).

Alpha-CENTAURI (Sevim et al., 2016) addresses a problem similar to the String Decomposition Problem. It takes a set of long error-prone reads as an input and uses them to generate the most likely set of monomers. It further decomposes each read into such monomers. However, Alpha-CENTAURI does not use additional information about previously inferred HOR structures and has a rather high rate of the incorrectly called monomers in the read decomposition.

Alpha-CENTAURI uses a pre-trained Hidden Markov Model (HMM) for a *consensus monomer* (consensus of all monomers over all alpha-satellite monomer families in the human genome). It aligns this HMM to all reads using the HMMer tool (Eddy, 1998) and clusters the generated alignments in order to identify monomer classes. Since this clustering is imperfect, the accuracy of string decomposition deteriorates as the tool reports spurious abnormal HOR structures (see Appendix “<u>Benchmarking string decomposition tools</u>”). However, the HMM alignment stage provides an accurate approximation for starting and ending positions of each monomer. Using the input set of monomers one can identify a monomer with the highest identity for each pair of these starting and ending positions, generate non-overlapping monomer alignments for each read, and transform each read into the monomer alphabet as described in the subsection “Transformation from the nucleotide alphabet to the block alphabet”. We refer to this approach (based on running HMMer on the consensus HMMs derived by Alpha-CENTAURI) as *AC*.

## Results

### Dataset

We analyzed the rel2 dataset of Oxford Nanopore reads (https://github.com/nanopore-wgs-consortium/CHM13) generated by the Telomere2Telomere consortium and released on March 2, 2019 (Miga et al., 2019). The dataset contains 11,069,717 reads (155 Gb total length, 50x coverage, the N50 read length equal to 70 kb) generated from the CHM13hTERT female haploid cell line. We used the rel2 base-calling with Guppy Flip-Flop 2.3.1. This read-set includes 999,562 *ultralong* reads (longer than 50 kb) that have the biggest impact on the centromere assembly and result in ≈32x coverage of the human genome.

We benchmarked various approaches to string decomposition using centromeric reads from chromosome X since this centromere (referred to as cenX) was recently assembled, thus providing the ground truth for our benchmarking. This benchmarking utilized 2,680 reads (total read length 132,9 Mb) that were recruited to cenX in Bzikadze and Pevzner, 2019).

### monomers and HOR sequences on cenX

We used the cenX HOR consensus sequence DXZ1* derived in Bzikadze and Pevzner, 2019. Appendix “<u>Extracting monomers from DXZ1*</u>” describes decomposition of DXZ1* into twelve monomers using Alpha-CENTAURI (Sevim et al., 2016). We denote these twelve monomers by letters from A to L and denote the consensus HOR sequence as ABCDEFGHIJKL (See Appendix “<u>cenX monomers</u>”).

### Reference cenX sequence

We analyzed the cenX v0.7 sequence assembled in Miga et al., 2019. 1,442 out of all 2,680 cenX reads were aligned to the reference using tandemMapper (Mikheenko et al., 2019). These reads form the set of mapped reads with total length 121 Mbp and with total length of aligned fragments 76 Mbp (see Appendix “<u>Generating accurate alignments</u>”).

Since each mapped read is typically aligned over its substring (rather than the entire read), it can be represented as a concatenate of a non-aligned prefix, an aligned substring, and a non-aligned suffix. We will find it convenient to trim the non-aligned prefix and suffix in each read, resulting in shorter reads forming a read-set *Reads*. Each shortened read *Read* is now aligned to a substring in cenX that we refer to as *origin(Read)*.

### Translating centromere and centromeric reads into monomer alphabet

Since each read *Read* in *Reads* is aligned to a substring *origin(Read)* in cenX, we can compare the sequence *mono(Read)* with the accurate sequence *mono(origin(Read))* representing the “ground truth” with respect to transforming sequences in the monomer alphabet. Using this approach, we benchmark the SD and AC approaches.

We used the SD tool to transform the cenX sequence (3.1 Mb) into the *mono(*cenX) sequence consisting of 18103 reliable monomers and 36 gap symbols (“?”) in the cenX region occupied by the LINE repeat. This conversion is reliable since monomers are rather conserved across cenX (median percent identity 98.8%). Figure 2 presents the distribution of percent identities of twelve cenX monomers and the gap monomers.

**Figure 2.**
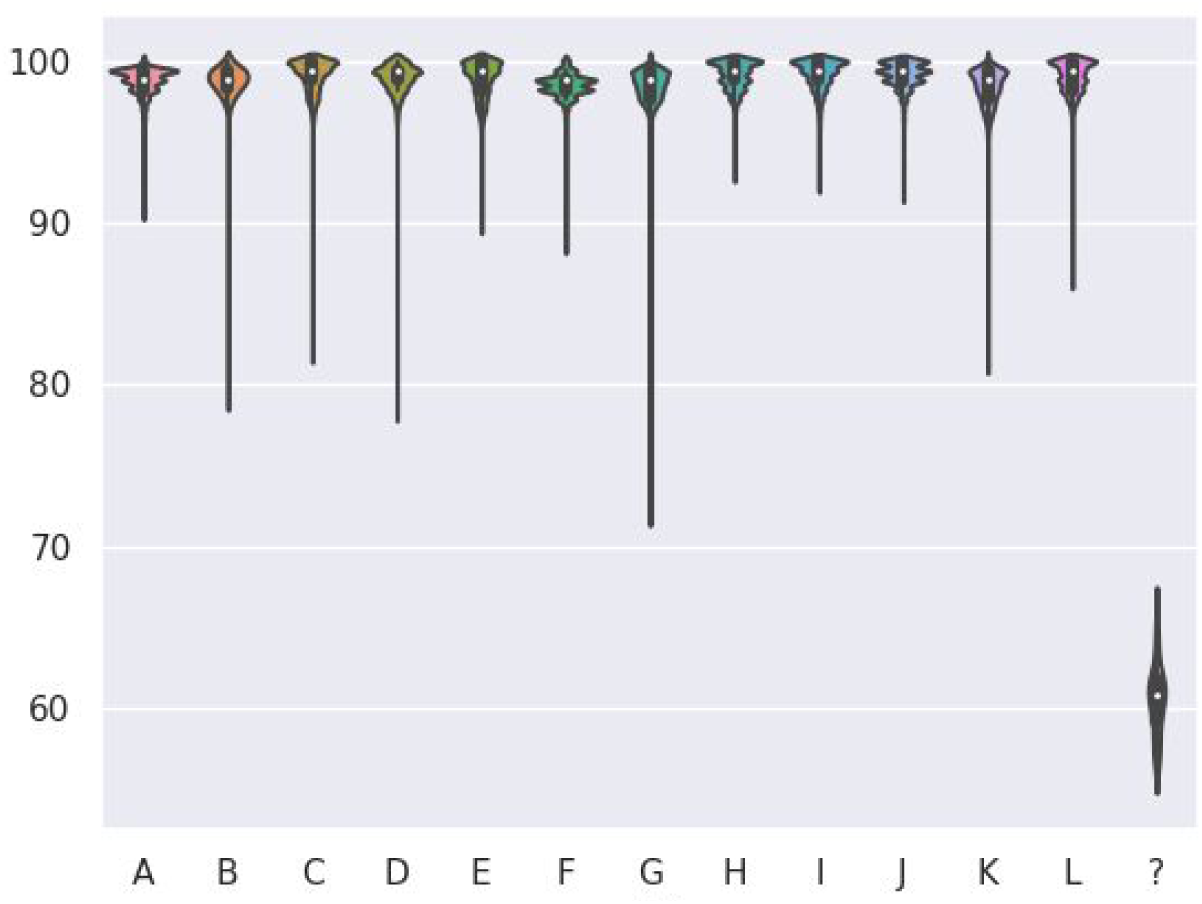
Distribution of percent identities of twelve cenX monomers and the gap monomer “?”. Each violin plot represents the distribution of the percent identity of a particular monomer across cenX. A LINE element at positions 2773652-2779726 is represented by 36 consecutive “?” symbols in the monocentromere X.

### Monoread-to-monocentromere alignments

We launched AC and SD to transform each read in *Reads* into a *monoread mono*(*Read)* and aligned it against *mono(origin(Read))* using the edit distance scoring (indel and mismatch penalties equal to 1 and match score equal to 0). Each alignment column contains a pair of symbols with the first symbol corresponding to a position in *mono*(*origin*(*Read*)) and the second symbol corresponding to a position in *mono*(*Read*). The symbols include twelve monomers, «?» symbol, and «-» symbol. We classify each column as a match or an error of one of the following types:

- monomer-monomer mismatch (monomer, monomer);
- monomer-gap mismatch (monomer, ?);
- monomer-deletion (monomer, -);
- gap-monomer mismatch (?, monomer);
- gap-deletion (?, -);
- monomer-insertion (-, monomer);
- gap-insertion (-,?);

Table 1 shows the error statistics for *Reads* and illustrates that AC generates four times more monomer-gap mismatches than SD (3414 vs 779). The monomer-gap mismatches usually occur in corrupted regions of reads, where the identity of the monomer-monomer matches flanking these regions usually falls below 80% (see Appendix “<u>Detailed analysis of errors in string decomposition</u>”).

**Table 1.** Summary of errors in the monoread-to-monocentromere alignments computed by the AC (black) and SD (blue) tools. Symbol “monomer” corresponds to one of the twelve cenX monomers, “?” corresponds to a gap symbol, “-” corresponds to a space symbol representing an indel in alignment of *mono(Read)* against *mono(origin(Read)).* A cell (*i, j*) represents the number of times when a symbol of type *i* in *mono(origin(Read))* was aligned to a symbol of type *j* in *mono(Read).* The number of matches is 445574 for SD and 442843 for AC.

| reads \ centromere | SD | AC | SD | AC | SD | AC | SD | AC |
| --- | --- | --- | --- | --- | --- | --- | --- | --- |
|  | monomer |  | ? |  | - |  | Total |  |
| monomer | 127 | 151 | 779 | 3414 | 115 | 157 | 1021 | 3722 |
| ? | 0 | 0 | - | - | 19 | 19 | 19 | 19 |
| - | 1 | 8 | 15 | 21 | - | - | 16 | 29 |
| Total | 128 | 159 | 794 | 3435 | 134 | 176 | 1056 (0.23%) | 3770 (0.84%) |

Overall, SD resulted in four-fold reduction in errors as compared to AC (779 versus 3414). It may appear that the AC tool is already accurate (0.84% error rate) and a reduction in the error rate (from 0.84% to 0.23%) is a useful but not critically important advance. However, it is crucially important since it provides information about much longer *k*-mers in monoreads thus enabling their assembly into a highly repetitive monocentromere (Bzikadze and Pevzner, 2019). For example, under an (unrealistic) assumption that errors are uniformly distributed, SD provides information about 434-mers, while AC provides information only about 121-mers in monoreads (100/0.16=625, while 100/0.86=116).

Below we analyze errors made by AC and SD in details.

### Analyzing monomer-monomer mismatches

104 out of 127 monomer-monomer mismatches made by the SD approach represent substitution of the monomer K by the monomer F (Figure 3). This substitution represents the most frequent mismatch for both approaches (115 out of 151 for AC). We explored the monoread alignments that substitute F by K and found that all such alignments correspond to the non-standard 16-monomer HOR ABCDEFGHIJ**F**GHIJKL, where the second occurence of F is often replaced by K in monoreads. A similar situation can be seen for K into L substitutions (3 and 5 mismatches for the SD and AC approaches respectively) in alignments of the non-standard 11-monomer HOR ABCDEFGHIJ**K**.

**Figure 3.**
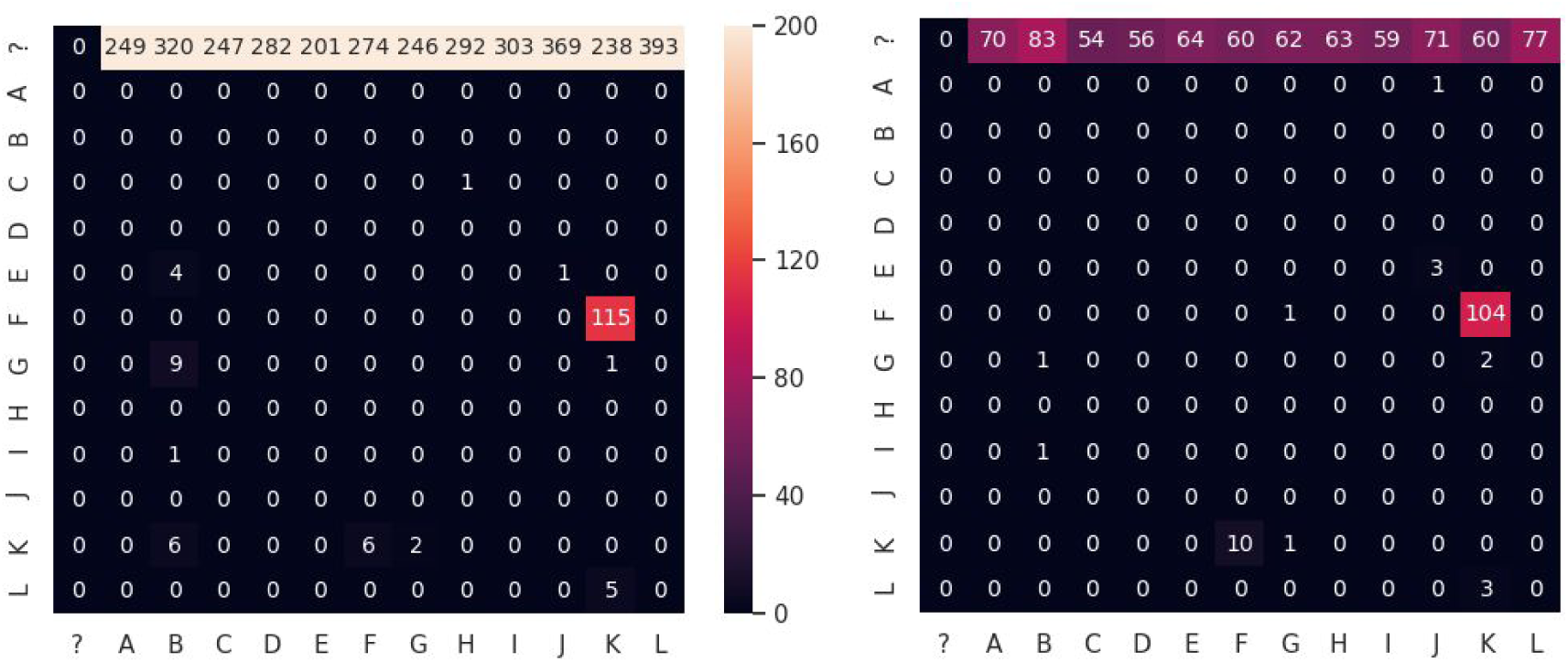
Mismatch substitution matrices for AC (left) and SD (right). The (X,Y) cell shows the number of times when symbol X in the monocentromere X (monomer or the gap symbol «?») was replaced by symbol Y in the monoread.

There are 12 occurrences of the non-standard HOR ABCDEFGHIJ**F**GHIJKL in the monocentromere X. We computed pairwise percent identities for nucleotides sequences of each of twelve occurrences of **F** in monocentromere and also compared them to monomers F and K (Figure 4, left).This analysis reveals two clusters: monomers **F** at HORs 1—4 and 5—12. Monomers 1, 3, 4 are identical and very close (98%) to monomer 2 and to the monomer F (median percent identity 98%). Thus, monomers 1-4 likely represent slightly diverged copies of F.

**Figure 4.**
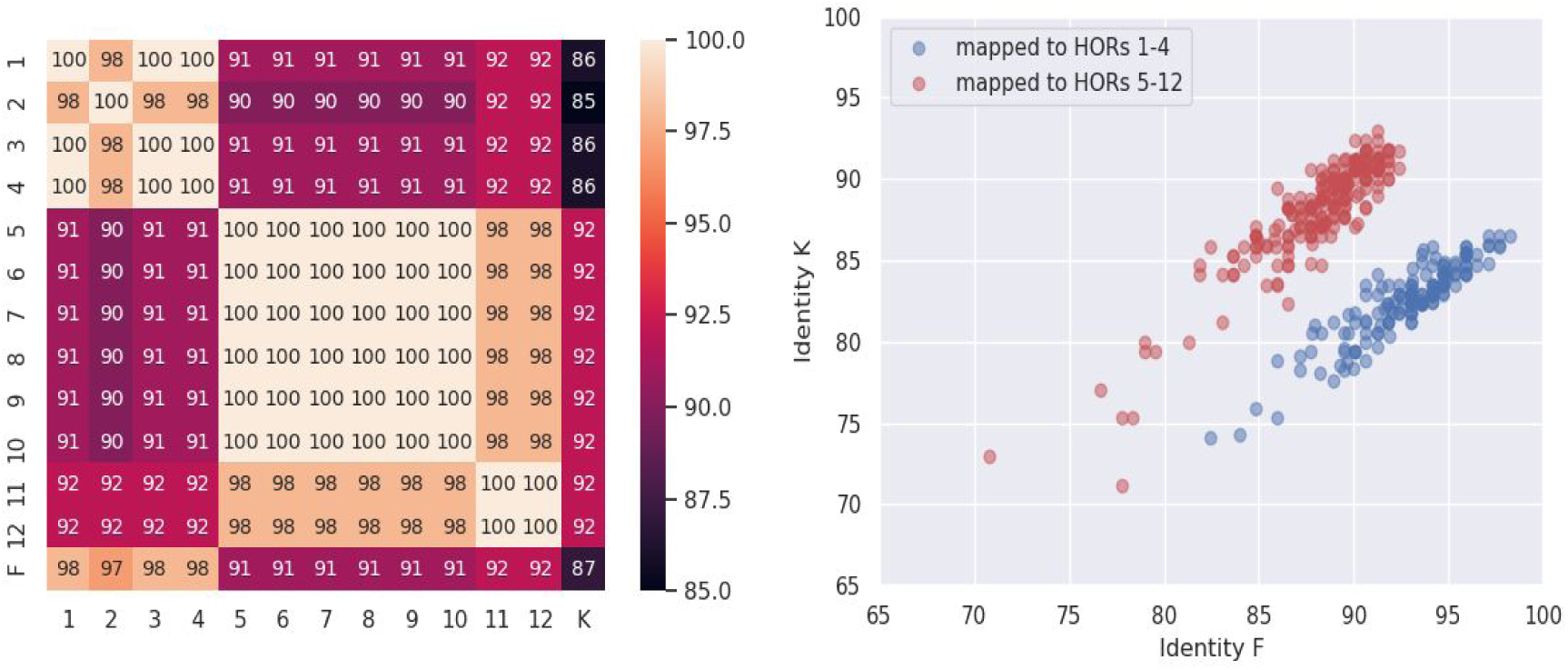
Analysis of the non-standard HOR ABCDEFGHIJFGHIJKL. (Left) The heatmap of identities between the occurrences of **F** in twelve non-standard HORs ABCDEFGHIJ**F**GHIJKL in cenX as well as monomers F and K. (Right) For all 377 occurrences of ABCDEFGHIJ**F**GHIJKL or ABCDEFGHIJ**K**GHIJKL in *Reads*, we computed the identity of 377 alignments of the monomer F (x-axis) and the monomer K (y-axis) against the corresponding substring of the read. Blue (red) circles represent occurrences of monomer F or K aligned to **F** from HORs 1—4 (HORs 5—12).

The second cluster consisting of **F** monomers at HORs 5—12 can be divided into subclusters of identical monomers 5—10 and 11—12 (similarity between monomers from these clusters is 98%). Surprisingly, monomers 5—12 **F** appear to be equidistant from both monomer F and monomer K with rather low sequence identity 91-92%. Analysis of pairwise alignments between these eight monomers and monomers F/K reveals that monomers 5-12 likely represent a chimeric monomer formed by the first half of K (first 81 positions of K) and second half of F (90 last positions of F), referred to as K+F (Figure 5). The monomer K+F has identity ∼97% with all of eight monomers in the second cluster.

**Figure 5.**
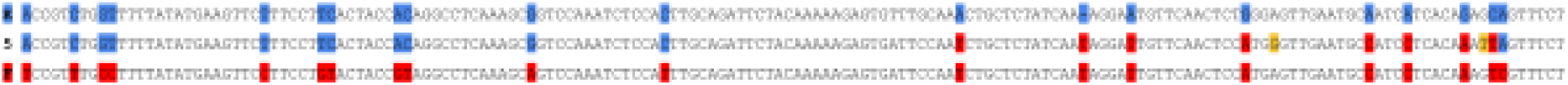
A 3-way alignment between the monomer F (1st row), monomer F from 5th instance of the non-standard HOR ABCDEFGHIJFGHIJKL (2nd row), and monomer K (3rd row). Positions that differ in F and K are colored by red in F and by blue in K. Positions of monomer **F** are colored either by red (if they have the same nucleotide as in F) or blue (if they have the same nucleotide as in K). Two positions in **F** that differ from positions in both F and K are colored in yellow.

The non-standard HOR ABCDEFGHIJ**F**GHIJKL was found in 272 monoreads generated by the SD approach (a similar HOR ABCDEFGHIJ**K**GHIJKL was found in 105 monoreads). All 272+105=377 occurrences of these HORs originated from the non-standard HOR ABCDEFGHIJ**F**GHIJKL in cenX. We classify these occurrences in two groups: 154 of them correspond to the first cluster (sequences 1—4 in Figure 4, right) and 377 - 154 = 223 correspond to the second cluster (sequences 5—12). For each of these 377 occurrences we extracted the nucleotide sequence that corresponds to **F/K** and aligned them to monomers F and K. Figure 4, right reveals two clusters of these 377 monomers that correlate well with grouping based on two clusters in Figure 4, left, confirming that a chimeric monomer K+F is supported by the reads.

We hypothesize a potential mechanism that generated the chimeric monomer K+F. Two cuts were introduced to the canonical 12-mers ABCDEFGHIJKL in the middle of monomers K and F resulting in trimmed sequences ABCDEFGHIJK’ and ‘FGHIJKL that were further glued together to form a non-standard 16-mer ABCDEFGHIJ(K+F)GHIJKL. Our identification of a novel K+F monomer suggests that a set of chromosome-specific monomers may quickly change during evolution by adding and potentially expanding chimeric monomers. It also emphasizes the importance of careful identification of all (even rare) monomers for follow-up centromere assembly efforts.

We decided to recompute monoread-to-monocentromere alignments with (K+F) monomer added to the original set of 12 monomers. We converted each read from *AlignedReads* and cenX sequence into 13-monomer alphabet. In the new representation monocentromere, the abnormal HORs 5—12 contain monomer K+F instead of **F**. The number of monomer-to-monomer mismatches for both approaches greatly decreased (from 127 to 17 for SD and from 151 to 47 for AC) and the number of other errors hardly changed (Table 2).

**Table 2.** Summary of errors in the monoread-to-monocentromere alignments computed by the AC (black) and SD (blue) tools for 13 monomers with additional (K+F) monomer. Symbol “monomer” corresponds to one of 13 monomers, “?” corresponds to a gap symbol, “-” corresponds to a space symbol representing an indel in alignment of *mono(Read)* against *mono(origin(Read)).* A cell (*i, j*) represents the number of times when a symbol of type *i* in *mono(origin(Read))* was aligned to a symbol of type *j* in *mono(Read).* The number of matches is 445684 for SD and 442946 for AC.

| reads | SD | AC | SD | AC | SD | AC | SD | AC |
| --- | --- | --- | --- | --- | --- | --- | --- | --- |
| centromere | monomer |  | ? |  | - |  | Total |  |
| monomer | 17 | 47 | 779 | 3414 | 115 | 158 | 911 | 3619 |
| ? | 0 | 0 | - | - | 19 | 19 | 19 | 19 |
| - | 1 | 6 | 15 | 21 | - | - | 16 | 27 |
| Total | 18 | 53 | 794 | 3435 | 134 | 177 | 946 (0.21%) | 3665 (0.81%) |

Non-standard 11-monomer ABCDEFGHIJ**K** has only one occurrence within monocentromere X and five occurrences in monoreads (a similar 11-monomer ABCDEFGHIJ**L** has three occurrences in monoreads). The monomer **K** in this 11-monomer has a rather low identity to the monomers K and L (87%). But it has identity ∼96% to a chimeric monomer K+L constructed from the first part of monomer K (the first 66 positions) and second part of monomer L (the last 105 positions). For all occurrences of 5+3=8 of non-standard 11-monomer in reads, the last monomer predicted as K or L has higher identity to K+L (∼90-94%) than to either monomer K (∼82-86%) or to monomer L (∼85%).

### Similarity between various monomers

Figure 6 illustrates that distinct monomers forming cenX have rather low similarity with each other (less than 80% for most monomer pairs). However, other HORs may contain more similar monomers that may lead to monomer-to-monomer substitution errors made by string decomposition algorithms.

**Figure 6.**
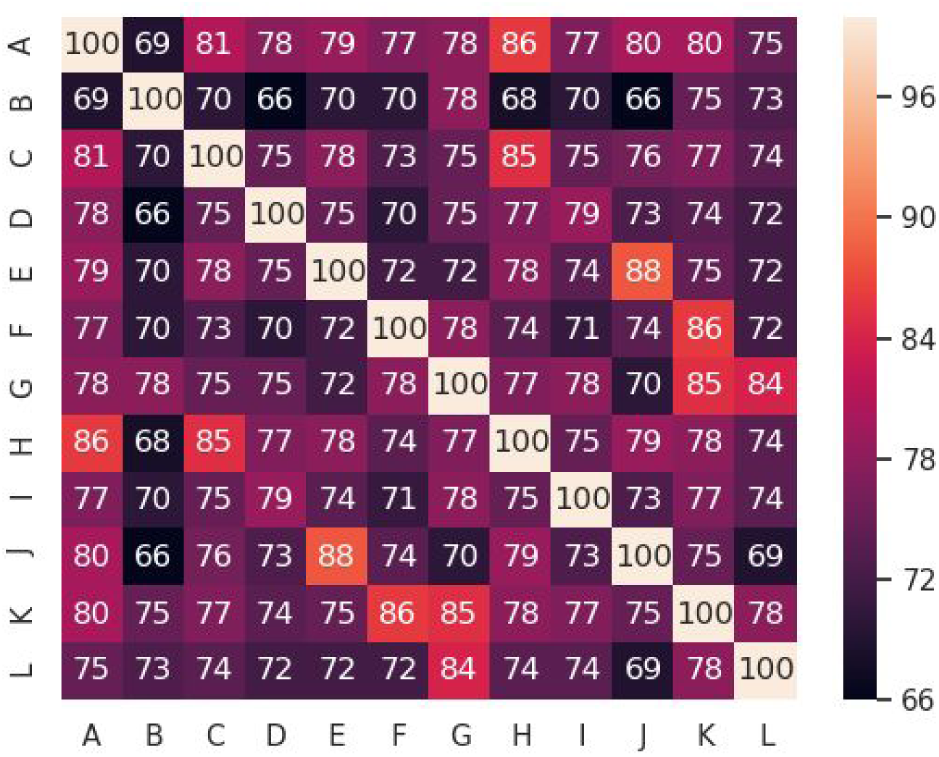
The monomer similarity matrix for twelve cenX monomers.

In order to analyze how string decomposition algorithms perform in the case of very similar monomers, we added a 13-th artificial monomer Z to the original set of 12 monomers. Monomer Z was constructed from monomer A by applying 3, 5, or 8 random nucleotide substitutions at randomly selected positions to the monomer A sequence (the resulting artificial monomers are referred as Z3, Z5 and Z8 respectively), Adding an extra monomer Z to the set of twelve monomers hardly affects the number of errors in Table 1, except for the number of monomer-monomer mismatches, when some monomer was substituted by Z. After adding the 13-th monomer, the number of monomer-monomer mismatches increased by 344/116/1 for Z3/Z5/Z8 for the SD approach (795/428/0 for the AC approach). This analysis illustrates that SD is better suited for string decomposition in the case when the monomer-set contains highly similar monomers.

## Discussion

StringDecomposer (SD) is the first tool designed specifically for decomposing long error-prone reads from ETRs (including nested ETRs such as centromeres) into blocks. We demonstrated that SD solves the String Decomposition Problem and accurately transforms long error-prone reads from centromeric regions into monoreads. Our simulations revealed that it remains highly accurate even in the case when the monomer-set contains highly similar monomers with percent identity as high as 98%. We thus project that SD will accelerate the ongoing centromere assembly efforts and will help to close the remaining gaps in the human and other genomes. It also promises to contribute to discovery of novel emerging (albeit still rare) monomers in the human genome as illustrated by our identification of the emerging K+F and K+L monomers on cenX.

## Code availability

StringDecomposition tool is publicly available on Github https://github.com/ablab/stringdecomposer. All scripts that were used for statistics calculation in Results section are available in https://github.com/TanyaDvorkina/sdpaper.

## Acknowledgements

We are grateful to Ivan Alexandrov, Karen Miga, and Alla Mikheenko for many useful insights. This work was supported by St. Petersburg State University, St. Petersburg, Russia (grant ID PURE 28396291).

## Appendices

- <u>SD implementation details</u>
- <u>Processing gaps in monomer alignment</u>
- <u>Benchmarking string decomposition tools</u>
- <u>Monomer-free benchmarking</u>
- <u>Extracting monomers from DXZ1*</u>
- <u>cenX monomers</u>
- <u>Generating accurate alignments</u>
- <u>Detailed analysis of errors in string decomposition</u>

## Appendix: SD implementation details

The implementation of the SD algorithm is publicly available on GitHub https://github.com/ablab/stringdecomposer. The main script “run_decomposer.py” accepts as input (i) a file containing reads or a genomic sequence, and (ii) a file containing monomer sequences (both in fasta format). It runs the SD algorithm (implemented in C++) and converting alignment scores of individual monomers into percent identities using Edlib library (Šošic and Šikic, 2017). Finally, it saves the monomer alignments to a given read-set (or genomic sequence) in the tsv-format. Below we highlight some implementation details.

### SD parallelization

The String Decomposition Graph may become rather large in the case when the string *R* is long (e.g., when *R* is an ultralong read or an assembly of an entire centromere) or when the block-set is large, leading to a large memory footprint. To reduce memory footprint, we represent the string *R* as a set of short overlapping segments of length *SegmentLength* with default value *SegmentLength =* 5500 bp (the last segment can be shorter) so that the consecutive segments overlap by *Overlap* nucleotide (Figure S1). The default value *Overlap* (500 bp) is chosen to be larger than the monomer size and to ensure that each monomer is positioned fully inside at least one segment. Afterwards, each segment of the read is processed separately and all segments are “glued” together.

**Figure S1.**
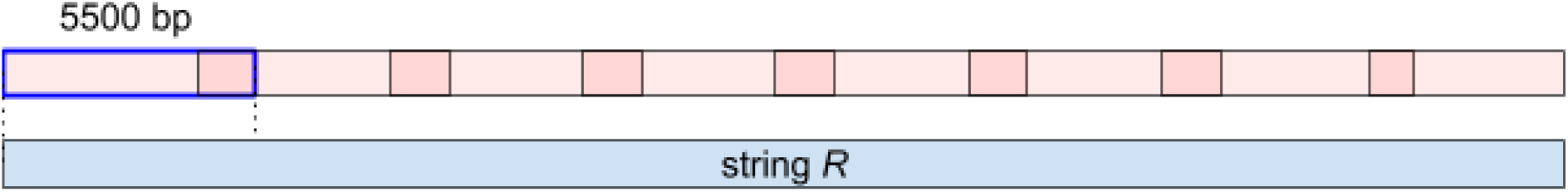
Partitioning reads into overlapping segments.

### SD parameters

The default penalties for indels and mismatches are equal to 1. However script “run_decomposer.py” allows to assign them arbitrary user-defined values.

### Selecting *MinIdentity* threshold

In order to select the *MinIdentity* threshold for identification of the gap symbols “?”, we generated a set of 100,000 random nucleotide sequences of size 171 bp, aligned each of them against each of 12 monomers from cenX, and selected one of these twelve monomers with the highest percent identity *Identity*. Although the average value of *Identity* across 100,000 random sequences is rather low (61%) some sequences resulted in much higher values, illustrating a potential risk of misidentifying a random sequence as a monomer (Table S1). Analysis of the distribution of identities (Table S1) suggests 69% as a reasonable choice of the *MinIdentity* threshold.

**Table S1.** Fraction of randomly generated sequences with *Identity* exceeding the given identity threshold *MinIdentity* (for *MinIdentity* varying from 60% to 70%).

| *MinIdentity* (%) | *fraction of random sequences exceeding MinIdentity* |
| --- | --- |
| 60 | 0.72939 |
| 61 | 0.47397 |
| 62 | 0.22885 |
| 63 | 0.14046 |
| 64 | 0.04128 |
| **65** | **0.00892** |
| 66 | 0.00354 |
| 67 | 0.00034 |
| 68 | 0.00008 |
| 69 | 0.00002 |
| 70 | 0.00002 |

Additionally we divided chromosome X assembly (Miga et al., 2019) into segments of length 5 kb and ran SD on these segments (with twelve cenX monomers) in order to analyze spurious alignments between non-centromeric regions and cenX monomers and to check if segments from chromosome X tend to be more similar to cenX monomers than random sequences. Using SD we built the chromosome X decomposition into monomers, Figure S2 shows the monomer alignment identity depending on its position on chromosome X. Most alignments outside the centromeric region have median identity ∼60%. The identity increases to ∼80% in the near-centromeric regions, and to 95% in the centromere, having a single draw-down in the cenX region occupied by the LINE element (varying from 60 to 70%).

**Figure S2.**
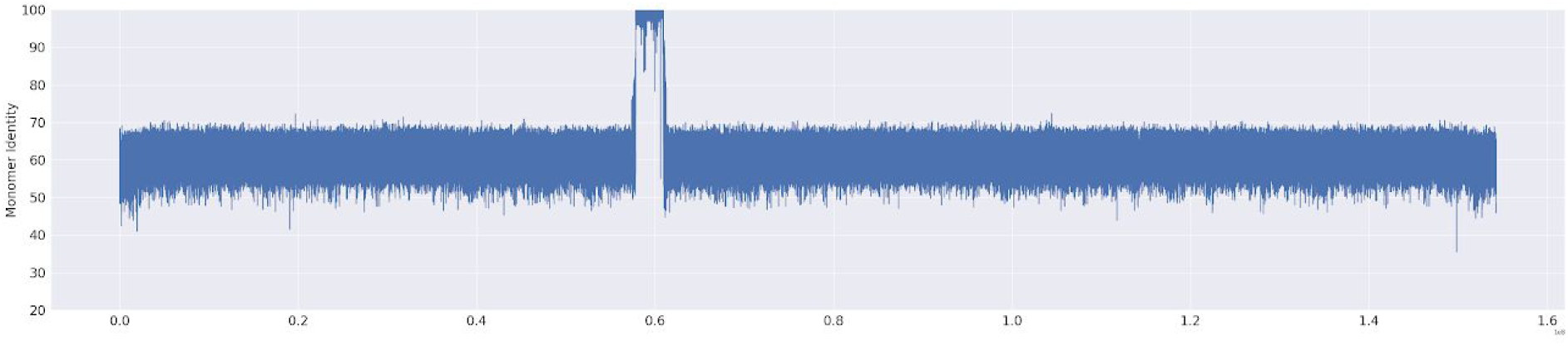
Monomer alignment identity across the entire chromosome X produced by the SD approach.

## Appendix: Processing gaps in monomer alignment

Some reads are translated into monoreads with gaps, where “?” symbols have low identity to all monomers. The SD algorithm generates a read decomposition where each position is covered by a monomer and replaces all unreliable monomers by the “?” symbol in the monoread. However, the number of predicted “?” in a read is not necessarily an accurate approximation of the total length of non-monomeric positions in a read. Additionally, the AC algorithm produces a decomposition that does not cover all positions of a read, resulting in *non-covered* positions in the AC monomer decomposition, with no monomer alignment covering these positions.

We thus modified a transformation of a read *R* into a monoread *mono(R)* by replacing a run of non-covered positions of length *L* by a run of the gap symbol “?” with length *L/MonomerLength*, where *MonomerLength* is the average length of monomers.

## Appendix: Benchmarking string decomposition tools

### Alpha-CENTAURI

Alpha-CENTAURI v.0.2 was run with default parameters. While HMMer search from the first stage of the Alpha-CENTAURI algorithm (partitioning a read into consequent monomers locations) was successful, the second stage (monomer sequences clustering and monomer identification) did not generate a precise read decomposition into monomers, reporting many abnormal HORs alignments, and was removed from further analysis.

### TandemRepeatsFinder

TRF 4.09 version was run with recommended parameters for human genome (https://tandem.bu.edu/trf/trf.whatnew.html). Though TRF has successfully identified the monomer length (∼170 bp) in 1926 reads, its output is difficult to use for further analysis. In particular, it is not clear how to identify monomers from the putative positions identified by TRFs as these positions are often shifted. For example a read *bcc5e5d2-f12f-4b59-b952-bd10f81ac89f* in rel2 T2T dataset is fully covered by DXZ1* monomers both according to SD and TRF, but TRF alignments positions have 40 bp shift regarding SD positions and monomers can not be identified with high identity scores. In contrast, the shift is rather small (∼10-15 bp) in a read *c500a3b1-f00c-40c1-94af-e33bae40ca71* resulting in a successful prediction of monomers from TRF alignments.

### NCRF

We launched the latest release of NCRF v1.01.02 to search for repeats of DXZ1* sequence with parameters “--scoring=nanopore --minlength=5000”. Appendix “Monomer-free metrics” reports NCRF results and compares it with other string decomposition approaches.

## Appendix: Monomer-free benchmarking

We compared NCRF with the AC and SD approaches using the dataset *Reads* defined in the Results section and analyzing two monomer-free metrics:

- *read coverage*, the fraction of reads’ length partitioned into monomers for (AC and SD approaches) or covered by the DXZ1* repeat (NCRF approach).
- *percentage of unaligned segments.* Two consecutive aligned monomers in a read are separated by an unaligned segment if the distance from the end of the first monomer alignment to the start of the second monomer alignment exceeds *MinUnalignedLength* (default value *MinUnalignedLength* = 10 bp). NCRF reports HOR without spaces in alignment, so it has 0 unaligned segments by definition.

Table S2 illustrates that NCRF has much lower *read coverage* the SD and AC approaches but by design improves on the AC and SD approaches with respect to the number of unaligned segments.

**Table S2.** Monomer-free metrics for AC, NCRF and SD.

| Approach | read coverage (%) | % unaligned segments<br>(#unaligned segments / #monomers) |
| --- | --- | --- |
| AC | 98 | 2.95 |
| NCRF | 92 | 0 |
| SD | 99 | 0.14 |

## Appendix: Extracting monomers from DXZ1*

In order to extract twelve monomer sequences we run “chop_to_monomers.py” script from Alpha-CENTAURI v.0.2G on the consensus monomer HMM (https://github.com/volkansevim/alpha-CENTAURI/blob/master/example/alpha.hmm) and a concatenation of two DXZ1* sequences derived in Bzikadze and Pevzner, 2019. The twelve cenX monomers are derived from the Alpha-CENTAURI output.

## Appendix: cenX monomers

The twelve monomers forming cenX HOR monomers are usually reported as CDEABCDEABCD since this sequence of monomers reflects the ancestral pentamer structure (CDEAB) of the HOR from which cenX HOR (DXZ1) originated. Since this representation is inconvenient for analyzing string decomposition of cenX, we represent DXZ1 as ABCDEFGHIKL instead.

## Appendix: Generating accurate alignments

In order to generate a set of accurate alignments, positions of alignments generated by tandemMapper were compared to the positions of read alignments in cenX assembly generated by centroFlye. It turned out that some tandemMapper alignments differ from centroFlye alignments. This likely caused either by incorrect read-to-centromere mapping (generated by tandemMapper) or erroneous recruitment of non-cenX reads to cenX (provided by centroFlye). We thus filtered out reads with differing starting positions (by more than 2 kbp) of tandemMapper and centroFlye alignments, resulting in 1442 read alignments.

## Appendix: Detailed analysis of errors in string decomposition

Most alignment errors between monoreads and monocentromere for both SD and AC approaches occur due to inconsistencies between (inaccurate) reads and (accurate) centromere assembly.

Since 91% of mismatches for the AC approach are *monomer-gap mismatches* (Table 1), we analyzed monomers predicted by SD but missed by AC. All SD monomer alignments that have overlap longer than 100 bp with some gap symbol (“?”) output by AC were considered. All monomer predicted by SD were divided into three groups: (i) the highest-scoring monomer is a true monomer, (ii) the second highest-scoring monomer is a true monomer, and (iii) none of the two highest-scoring monomers is a true monomer. Figure S3 presents the scatter-plot of the scores of the highest-scoring and the second highest-scoring monomers for each group (left) and the distribution of their differences (right). All alignments have relatively low identity (below 85%) as compared to the average identity of all monomers (93%). However, the highest-scoring monomer is correct in ∼99% of cases and the difference in identity between the highest-scoring and the second-highest scoring monomers is rather substantial (more than 4% in most cases). Both the highest-scoring and the second-highest scoring monomers are incorrect in approximately 0.3% of cases.

**Figure S3.**
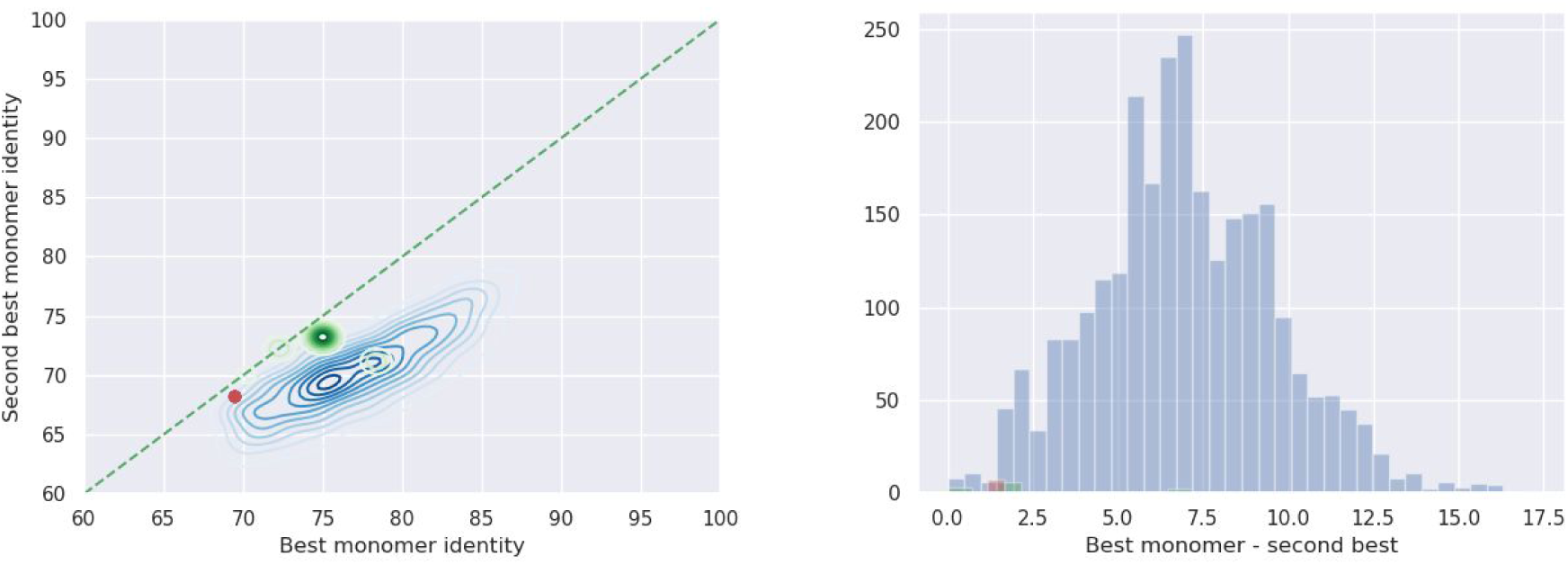
Statistics of scores for monomers that the AC approach failed to predict. (Left) Analysis of underpredicted monomer that were classified as monomer-gap mismatches: the scatter-plot of the highest monomer score and the second-highest monomer score for three cases: the highest-scoring monomer is correct (blue), the second highest-scoring monomer is correct (green), neither the highest-scoring nor the second highest-scoring monomer are correct (red). Intensity of the color reflects number of points with such identity values. (Right) Distribution of differences between the identity of the highest-scoring and the second highest-scoring monomers.

SD and AC made 15 (21) gap-insertions, 1(8) monomer-insertions, and 115 (157) monomer-deletions. Most such errors arise in corrupted regions of reads with low alignment quality — the identities of flanking monomers located next to such region usually falls below 80%. AC has more insertions (deletions) than SD, as the run of “?” identified by AC are sometimes longer (shorter) than the correct number of monomers in the run.

## Notes

https://github.com/ablab/stringdecomposer

